# Hyperreflexia after corticospinal tract lesion reflects 1A afferent circuit changes not increased KCC2 hyperexcitability

**DOI:** 10.1101/2025.02.21.639555

**Authors:** Thelma Bethea, Temitope Adegbenro, John H. Martin

## Abstract

Hyperreflexia is a consequence of spinal cord injury (SCI) and motor system lesions in the brain Two major mechanisms underpinning hyperreflexia have been reported: proprioceptive afferent (PA) circuit changes produced by 1A fiber sprouting, which could enhance reflex signaling, together with reduced GABAergic inhibitory presynaptic regulation (GABApre); and increased intrinsic motor neuron excitability, for example, produced by reduced motor neuron membrane-bound potassium-chloride co-transporter2 (KCC2). Here we examine how selective unilateral CST injury in the medullary pyramid (PTX), which eliminates the CST from one hemisphere, allows for specific investigation of the different mechanisms to determine their contributions to hyperreflexia. We used rate-dependent depression (RDD) of the Hoffmann (H)-reflex for the forelimb and hindlimb 5^th^-digit abductor muscles to assess hyperreflexia on both the contra-and ipsilesional sides. We compared RDD in naive and unilateral-PTX rats at 7-dpi and 42-dpi, supplemented with additional timepoints to examine hyperreflexia development.

Immunohistochemistry was used to identify PA synapses (VGlut1), GABA presynaptic boutons (GABApre), motor neurons (ChAT), and to measure KCC2. Following unilateral PTX, we observed significant hyperreflexia in the contralesional forelimb only. Membrane-bound KCC2 was unchanged in contralesional cervical motor neurons. Whereas both cervical and lumbar motor neurons showed increased PA sprouting contralesionally, there was a concomitant increase in GABApre terminals for the lumbar not cervical cord, which associated with a normal hindlimb H-reflex. Our findings show that KCC2 is disassociated from hyperreflexia in the uniPTX model. Instead, forelimb hyperreflexia can be explained by cervical motor neuron PA sprouting and an uncompensated GABApre regulation.

## Introduction

Hyperreflexia after motor cortex stroke or spinal cord injury (SCI) contributes to significant additional disease burden in patients suffering a loss, of skilled voluntary movements (Hiersemenzel et al., 2000; Hultborn, 2003). Two classes of mechanisms underpinning hyperreflexia have been identified. The first is increased intrinsic motor neuron excitability. For example, a reduction in the potassium-chloride co-transporter 2 (KCC2), a membrane-bound protein that maintains low intracellular [Cl^-^] with extrusion of K^+^ (Lee et al., 2007), occurs after SCI (Boulenguez et al., 2010). Increased developmental expression of KCC2 is associated with GABA-A and Glycine receptor mediated hyperpolarization (Boulenguez et al., 2010; Hübner et al., 2001). SCI produces a reduction in the membrane-bound KCC2, thereby driving up motor neuron excitability (Chamma et al., 2012), contributing to hyperreflexia after SCI (Boulenguez et al., 2010; Bilchak et al., 2021; Beverungen et al., 2020).

The second class of mechanism is enhanced signaling in the proprioceptive afferent (PA) circuit. SCI damages segmental afferent connections and intrinsic interneurons and circuits at the level of the injury, in addition to many sensory and motor projection systems (Oudega and Perez, 2012). Earlier we showed that unilateral selective elimination of the corticospinal tract (CST) from one hemisphere, by pyramidal tract lesion (PTX) in the caudal medulla, is sufficient to produce contralesional forelimb hyperreflexia (Tan et al., 2012). We further showed that this led to 1A afferent fiber sprouting on the affected side of the cervical spinal cord (Tan et al., 2012).

This increase in 1A synapses on motor neurons, which are VGlut1-positive, could contribute to hyperreflexia through enhanced 1A signaling. Subsequently, we showed that motor cortex inactivation, not injury, increased contralateral 1A afferent projections, suggesting that an imbalance in activity-dependent completion exerted by the silenced CST terminals is an important factor driving 1A afferent sprouting (Jiang et al., 2019; Jiang et al., 2016). Bilateral PTX also produces 1A afferent sprouting and, like SCI, a reduction in membrane-bound KCC2 (Kathe et al., 2016). That study also showed that there was a reduction in GABApre contacts on 1A motor neuron terminals.

The present study aimed to distinguish the potentially complementary contributions to hyperreflexia of KCC2-mediated motor neuron hyperexcitability and 1A circuit mechanisms (1A sprouting and associated GABApre inhibition). We used a unilateral PTX to cut all corticospinal projections from one hemisphere. These eliminates most CST axons in the contralateral spinal cord, sparing only the ipsilateral projections (2-5%). Hyperreflexia was assessed using rate-dependent depression of the H-reflex, (RDD). We characterized RDD changes for the forelimb and hind limb during an early subacute period (7 days post-injury) and the chronic period (42 days post-injury). Clinically, lower limb spasticity is less prominent (Sunnerhagen et al., 2019; Urban et al., 2010) and spasticity may improve function by promoting lower limb muscle tone (Dietz, 2003), but not so for the upper extremity (Kamper et al., 2003). We assayed the 1A connections on motor neurons and their presynaptic inhibition (GABApre) using highly sensitive motor neuron 3-D reconstruction, and segmental KCC2 measurement.

Our findings show contralateral forelimb but not hindlimb hyperreflexia following unilateral PTX. Although sprouting of 1A afferent terminals on motor neurons occurs in both the contralateral cervical and lumbar enlargements, GABApre sprouting commensurate with the 1A sprouting only occurs at the lumbar level, where hyperreflexia did not occur. Surprisingly, membrane-bound KCC2 did not change in either the cervical or lumbar enlargements, showing a 1A circuit-based, not intrinsic motor neuronal excitability, mechanism for hyperreflexia. Thus, using a model brain lesion that axotomizes all CST axons from one hemisphere, we show by exclusion that, among the three mechanisms considered, a lack of effective GABApre regulation explains hyperreflexia in this model.

## METHODS

### Animals

Experiments were performed in accordance with the National Institutes of Health Guidelines for the Care and Use of Laboratory Animals. All animal protocols were approved by the Advanced Science Research Center Institutional Animal Care and Use Committee. Adult male Sprague Dawley rats (250–275 g) were used for this study. Animals were housed under a 12 h light/dark cycle in a pathogen-free area with water and food provided ad libitum.

Animals comprised three principal groups: naïve (uninjured), early post-injury (7-days-post-injury, dpi), chronic post-injury (42-dpi).

### H-Reflex Recording

Animals were anesthetized with an induction dose of ketamine and xylazine (70mg/kg ketamine; 5 mg/kg Xylazine, i.p.) and maintained on ketamine alone during the course of physiological experiments (Hosoido et al., 2009). Animals were monitored continuously throughout all procedures and supplemental doses of ketamine (25mg/kg) were provided as needed. To determine the rate-dependent depression (RDD) of the H-reflex (Ho and Waite, 2002; Tan et al., 2012), paired-pulse stimulation with control and test pulses were used at increasing inter-pulse intervals (IPIs) (20-2000 ms). We used percutaneous muscle stimulation and recorded the evoked the direct M (muscle) and H-reflex responses through the same electrodes. For the forelimb, we studied the abductor digiti quinti muscle (ADQ) and, for the hindlimb, the abductor digiti minimi muscle (ADM). We chose the percutaneous method (Kathe et al., 2016) because we wished to examine the time-course of development of hyperreflexia repeatedly in the same animal (see Supp Fig1). In a prior study involving the extensor carpi radialis (ECR) muscle, we found that this percutaneous method did not reliably distinguish the M-and H-waves. In initial experiments, we found that ADQ and ADM H-reflex recordings using this method produces reliable separation of the M-and H-waves, resulting in consistent RDD. Thus, we were able to use percutaneous stimulation/recording repeatedly and consistently with this method.

### Retrograde labeling of motor neurons

Assessment of KCC2 was made in extensor carpi radialis (ECR) and tibialis anterior (TA) motor neurons of the cervical and lumbar segments. We confirmed the identities and locations of the motor neurons by intramuscular injection of cholera toxin subunit B (CTB; FITC conjugated, List Biological Laboratories) into the muscles, to retrogradely label the motor neurons. CTB injections were made 7 days prior to euthanizing the animals (7-dpi early time point group; 42-dpi later timepoint group), which is sufficient time for retrograde labeling of motor neurons and anterograde transport transganglionic labeling of 1A afferent terminals (Jiang et al., 2016). CTB (10 µm for each muscle, 1% solution) was injected into the target muscles bilaterally using a Hamilton syringe (Tan et al., 2012). A total of 40 μl of 1% CTB was injected into each rat.

### Unilateral Pyramidotomy

To perform the unilateral pyramidal tract injury, animals were anesthetized with Ketamine-Xylazine (70mg/kg Ketamine; 5 mg/kg Xylazine, i.p.). Supplemental doses of Ketamine (25 mg/kg) were administered as needed. Similar to our earlier studies (Jiang et al., 2016; Tan et al., 2012), an incision was made through the skin of the ventral neck. The tissues were bluntly dissected, and the trachea was displaced laterally, to expose the ventral surface of the occipital bone. Using a rongeur, a small craniotomy was made to expose the ventral medulla.

The dura was cut, using an iridectomy scissor (Fine Science Tools), to expose the pyramids. The scissor was used to cut the right pyramid, to a depth of 1.2 mm below the ventral medullary surface to completely transect the pyramid. Care was taken to not damage the basilar artery.

Completeness of the lesion was confirmed postmortem with histological assessment of the loss of PKC-γ immunoreactivity in the dorsal CST, contralateral to the injury, at C3.

### Tissue Processing and Immunohistochemistry

At the termination of experiments, rats were deeply anesthetized with a mixture of ketamine and xylazine (70/5 mg/kg, i.p.) and transcardially perfused with 250 ml of 0.1M phosphate buffer (PB) followed by 500ml of 4% paraformaldehyde solution. The brain and spinal cord were removed and postfixed in paraformaldehyde solution for 3hrs before being transferred to a 30% sucrose solution. Tissue was sectioned transversely at a thickness of 40 μms. To identify 1A terminals on motor neurons and their presynaptic contacts, sections were labelled with: 1) the vesicular glutamate transporter 1(rabbit anti-Vglut1)(Millipore 1:5000 conc), a marker for PA terminals (Alvarez et al., 2004), 2) glutamic acid decarboxylase 65 (chicken anti-GAD65)(Millipore, 1:1000 conc.), a marker of GABAergic presynaptic inhibition, and 3) CTB (List Biologicals goat anti-CTB 1:2000 conc.), to label motor neurons. Secondary antibodies used included: guinea pig-Cy5 for Vglut1(Millipore 1:5000 conc.), rabbit-Cy3 for GAD65 (Jackson Labs 1:1000 conc.), and goat-FITC for CTB (Invitrogen 1:2000). A separate set of tissue sections from both spinal segments, at acute and chronic timepoints were stained for KCC2 (Millipore, rabbit-KCC2; 1:100 conc.), and co-stained with choline acetyltransferase ChAT (Millipore Goat-ChAT 1:100 conc.) that is used as a marker for motor neurons. Secondary antibodies used were: guinea pig-Cy5 for KCC2 (Millipore 1:5000 conc.) and goat-FITC against ChAT (Invitrogen 1:100).

### 1A afferent terminals Image Analysis

Sections within the cervical (C6-C8) and lumbar (L1-L4) spinal cord were visualized and imaged digitally using a Zeiss LSM 880 confocal microscope. Individual motor neurons as well as their proximal dendrites ipsilateral and contralateral to the injury were imaged as a z-stack at 63x magnification with a 2.23μm optical step size with 50% overlap. Motor neurons (soma and dendrites) were 3D-rendered along with accompanying 1A proprioceptive afferent (PA) terminals and GABApre bouton numbers using Imaris Software (Oxford Instruments ver. 9.7.1). Quantitative analyses were also performed using Imaris. We previously used ImageJ to count presynaptic 1A afferent synaptic puncta on motoneurons (Tan et al., 2012; Sharif et al., 2021). However, the counting of contacts in a single focal plane revealed the limitation of the 2-D analysis. By comparison, Imaris allowed us to visualize all potential interactions (i.e., contacts) by rendering all components from a z-stack confocal file into 3D rendered objects. This method allows us to accurately and consistently measure presynaptic 1A excitatory terminals, as well as GABApre inhibitory terminals, to assess morphologically the 1A circuit hypothesis for hyperreflexia. We used a series of thresholding steps for each component starting with CTB-positive motor neurons, then Vglut1-positive synapses, then GAD65-positive synapses. Interactions between rendered motor neurons and Vglut1-positive synapses were defined by a separation distance of zero micrometers. Similarly, interactions between rendered Vglut1-positive synapses that contact the motor neuron, and rendered GAD65 synapses were defined by a separation distance of zero micrometers. Motor neurons were imaged and analyzed individually in the cervical (ipsi-lesional: n=37 motoneurons; contralesional: n=33 motoneurons) and lumbar (ipsilesional: n=20 motoneurons; contralesional: n=20 motoneurons) segments for the early timepoint. For the late timepoint, the cervical (ipsilesional: n=53 motoneurons; contralesional: n=50 motoneurons) and lumbar segments (ipsilesional: n=39 motoneurons; contralesional: n=46 motoneurons).

### KCC2/ChAT Image Analysis

KCC2 in motor neurons, identified using ChAT immuno-staining, within cervical (C6-C8), and lumbar (L1-L4) segments were analyzed (ImageJ analysis software, ver. 1.53a). Three lines were drawn cross-sectionally through each motor neuron to measure KCC2 labeling grayscale pixel intensity at the two membrane intersection points and in the cytosol.

Supplemental Figure 1 shows a ChAT/KCC2 double-labeled motor neuron (A) and one representative line for constructing the average histogram (B). Histograms were generated for each line. Area under the curve for the membrane, and cytosolic regions were measured as integrated pixel intensities. The intensity for the membrane and cytosolic regions were averaged (Kathe et al., 2016), and a ratio of membrane bound over cytosolic bound KCC2 was determined (Boulenguez et al., 2010). The mean intensity of membrane bound and cytosolic bound KCC2 were separately determined by normalizing gray matter KCC2 labelled motor neurons, to white matter KCC2 labelling (Boulenguez et al., 2010). The following motor neurons were analyzed: cervical (naïve right=19, left=20; early ipsilesional: n=25; early contralesional: n=26; chronic ipsilesional: n=17; chronic contralesional n=19), and lumbar (naïve right=24, left=22; early ipsilesional: n=21; early contralesional: n=23; chronic ipsilesional: n=19; chronic contralesional n=19).

### Western blots

To validate immunohistochemical KCC2 measurements, we used western blotting.

Animals were euthanized at 7 days post unilateral CST injury. The cervical (C4-C6) and lumbar (L3-L5) enlargements of the spinal cord (Naïve N=5; PTX N=3) were dissected, split along the midline to separate ipsilesional and contralesional segments, and placed into ice cold Hank’s balanced salt solution (HBBS; Invitrogen). The segments were placed in 4 microtubes and homogenized promptly in N-PER buffer (Invitrogen) supplemented with a protease-inhibiting cocktail (Invitrogen). Approximately 25 μg of protein was subjected to SDS-acrylamide-bisacrylamide gel electrophoreses along with a protein ladder (Crystalgen) for size comparison. Following electrophoresis, the protein bands were transferred to PVDF membrane (Millipore) and probed against the primary antibodies. An ECL chemiluminescent system (Cell Signaling Technology) was used to develop the Western blots. The following primary antibodies were used: rabbit anti-KCC2 (Millipore) at 1:500; and mouse anti-GAPDH at1:5000 (Sigma Aldrich). Secondary antibodies used were: guinea pig for KCC2 (1:10 conc.; Millipore) and goat for GAPDH (1:1000 conc.; Vector Laboratories).

### Statistics

Statistical analysis was performed using GraphPad (Prism Software, ver. 9.1.2). Unpaired Student’s t-test and one-way ANOVA followed by post hoc corrections for multiple comparisons were used as needed and described with results. Results are displayed as mean ± SEM, and P-value. Power analysis (G*power software, ver. 3.1.9.6) was used to guide animal numbers.

Estimates based on expected effect sizes for the lab’s prior plasticity and injury studies (effect size: f=5.7, a=0.05; b=0.85), are N=5/group for electrophysiology data and N=4 for immunohistology data. Analyses were performed by laboratory personnel blinded to the experimental protocol.

## Results

### Experimental design

We examined the physiological and morphological effects of a complete unilateral CST injury of the pyramid in the caudal medulla (PTX) at early (7-days post injury, dpi) and chronic (42-dpi) timepoints (Figure 1A). Unilateral PTX was verified to be complete in all animals by the absence of Pkc-γ staining at the C7 level (Fig 1B). H-reflex assessments (n=6 rats) were made prior to PTX (baseline) and at the early and chronic timepoints (Figure 1A). Additional timepoints were assessed (n=5 rats) to show the progression of hyperreflexia. H-reflex changes were assessed using the method of pair-pulse stimulation to measure rate-dependent depression (RDD) of the reflex (Fig 1C). Stimulation evokes the M-wave and the H-wave (Fig 1C). In a naïve animal (Fig 1C), the amplitude of the H-wave evoked by the second test pulse becomes reduced with short interstimulus intervals (ISI; Fig 1C, 2000 ms versus 20 ms). This is rate-dependent depression (RDD) of the H-reflex (Ho and Waite, 2002). Seven-days prior to euthanizing the animals, they were subjected to an intramuscular CTB injection into ECR and TA (see Methods) to localize their motor neurons for 1A afferent, and GABApre measurements.

**Figure 1.**
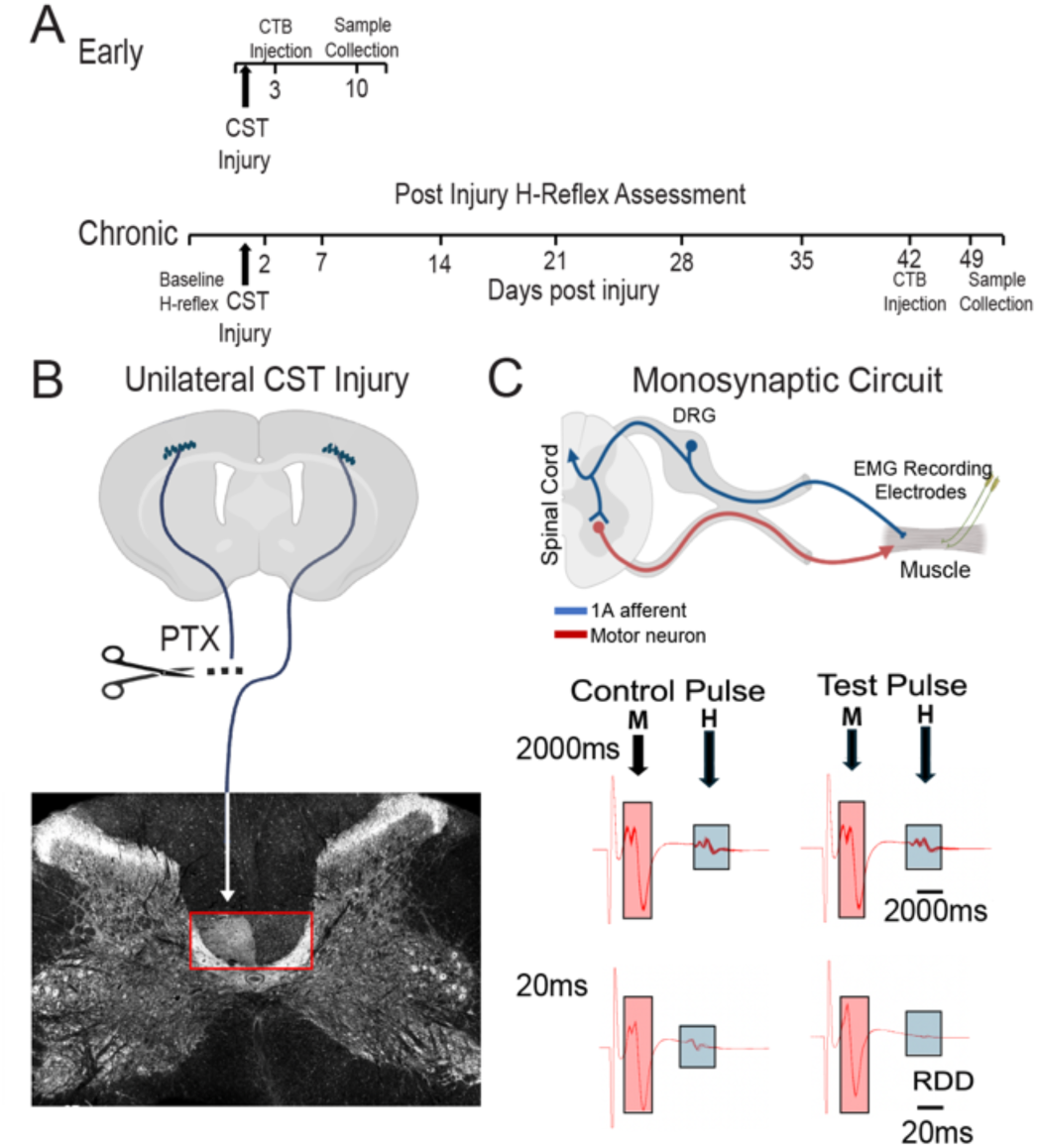
Experimental design. A. Time line of experiments. Early. Experiments were designed for collecting tissue samples only (CTB-labeled motor neurons, immunohistochemistry for KCC2, VGlut1, PKCγ and Gad65 (n=5 rats PTX; n=5controls/ shams). Chronic. Experiments designed for repetitive H-reflex testing (42-dpi) and collection of tissue samples for immunohistochemistry, as for the early cohort of rats (n=5). B. Unilateral PTX with PKCγ anatomical validation of lesion completeness. C. Rate dependent depression of the H-reflex. Top part of figure shows experimental setup with single electrode pair in the muscle to electrically activate afferent fibers and record the H-and M-responses.

These muscles were chosen because their motor pools are large and extend throughout most of the cervical enlargement and lumbar segments (Romanes, 1964; Hardman and Brown, 1985), making them good candidates for identifying a generalized motor neuronal response to CST loss. Additionally, ECR has been shown to display RDD after unilateral PTX (Tan et al., 2012). These animals were euthanized at 42-dpi, (chronic time point) and spinal tissue was collected for performing analyses to assess KCC2 and 1A terminals, together with GABApre terminals, on motor neurons. As indicated in the timeline, a separate set of animals we examined at 7-dpi (early time point; n=5) to assess morphology (KCC2 and 1A terminals) after PTX. Comparisons were made with naïve rats (n=5).

### Unilateral PTX produces forelimb, not hind limb, hyperreflexia

We first determined if there were differential effects of a unilateral CST injury on RDD of the H-reflex of the fore-and hind limbs. RDD plots (Figure 2A, B) show that the test H-wave changes as a function of the preceding conditioning pulse for the different ISIs. The plots for both ipsilesional limbs show the characteristic H-wave RDD reduction with progressively shorter ISIs and robust negative slopes (A,B, insets). By contrast, the contralesional forelimb shows that H-wave RDD at 100 ms, 50 ms, and 20 ms at both 1 week and 6 weeks is maintained; this indicates a reduction in RDD, our measure of hyperreflexia. Surprisingly, the contralesional hindlimb did not show any H-wave RDD reduction, and displays the characteristic negative slope. H-wave RDD reduction for the different conditions is compared at the 20 ms ISI (Fig 2C). Only the contralesional forelimb shows a significate reduction in RDD (i.e., larger test H-wave relative to the conditioning H-wave) at 7 dpi (one-way ANOVA; F=5.77; p=0.01) and 42 dpi (one-way ANOVA; F=5.77; p=0.02) compared to prePTX; all others are not significantly different (see Fig 2 legend for additional statistics). This indicates that hyperreflexia is present only for the contralateral forelimb. Importantly, that RDD is not different at the two post-PTX timepoints indicates that it also is persistent. We combined data from the H-reflex animals with that from an additional cohort (n=5), where we measured RDD after two additional post-injury time points (2-dpi and 14-dpi), in addition to 7-and 42 days, to determine the time course of development of hyperreflexia (Supplemental Figure 1). We observed a progressively larger test H-wave relative to the conditioning H-wave (i.e., RDD reduction; one-way ANOVA; F= 6.548), with a numerical RDD reduction at 2-dpi, significance from baseline by 7 dpi (p=0.005 Tukey post hoc), and maintenance of this difference at 42 dpi (p=0.0005). This suggests gradual development of hyperreflexia, as early as 2-dpi and maintenance from 6/7-dpi. Our findings show that a complete unilateral CST lesion results in only a segmentally specific forelimb hyperreflexia; is presents only in the contralesional forelimb and not the contralesional hindlimb.

**Figure 2.**
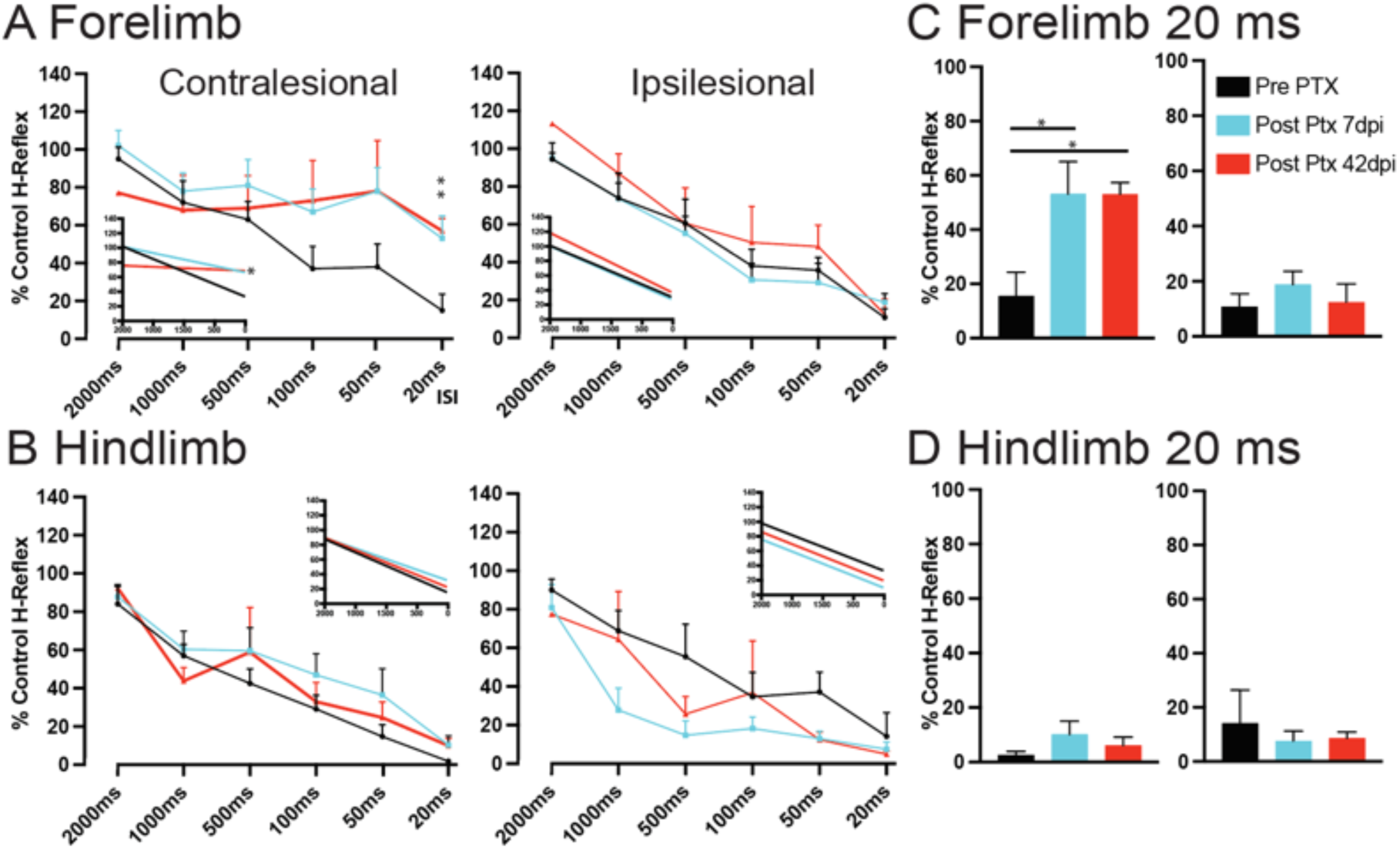
Rate-dependent depression. A, B. Rate-dependent depression plots for forelimb (A) and hindlimb (B). Insets show linear regression slopes for pre-PTX, post-PTX at 7-days, and post-PTX at 42-days. Color coding of groups is: pre-PTX, black; post-PTX at 7-days, light blue, post-PTX at 42 days, red (code also shown in part C). Linear regression plots for (A) Forelimb-Contralesional (slopes; Pre=-0.03435, 7 dpi=-0.01770, 42 dpi=-0.003551; F=7.065; p=0.009). (A) Forelimb-Ipsilesional (slopes; Pre=-0.03548; 7 dpi=-0.03673; 42 dpi=-0.04104; F=0.158; p=0.855). (B) Hindlimb-Contralesional (slopes; Pre=-0.03640; 7 dpi=-0.02919; 42 dpi=-0.03382; F=0.2397, p=0.7905). (B) Hindlimb-Ipsilesional (slopes; Pre=-0.03264; 7 dpi=-0.03326; 42 dpi=-0.03346; F=0.00419, p=0.9958). C, D. Rate-dependent depression measured at 20 ms for the forelimb (C, contralateral, left; ipsilateral, right; D, contralateral, left; ipsilateral, right). Contra-lesional hindlimb (Figure 2C) shows no significant difference from baseline at 7 DPI (one-way ANOVA; F=1.41; p=0.21) and 42 dpi (one-way ANOVA; F=1.41.77; p=0.48).

### Hyperreflexia is not due to KCC2-mediated intrinsic motor neuronal hyperexcitability

Reduction in membrane-bound KCC2, by internalization and subsequent cytosolic degradation after injury is a well-established mechanism supporting a role for motor neuron hyperexcitability in hyperreflexia (Boulenguez et al., 2010; Kathe et al., 2016; Cote et al., 2017; Bilchak et al., 2021). We next determined if unilateral PTX also produces a KCC2 reduction.

High-resolution confocal micrographs (Supplemental Fig 1A; Figure 3A) show characteristic dense KCC2 membrane staining in motor neurons in the naïve animals. Surprisingly, after PTX, we did not observe a change in KCC2 staining for the contralesional cervical or lumbar segments (Figure 3A). Recall that animals showed significant hyperreflexia only for the contralesional forelimb. We quantified the ratio of membrane-bound to cytosolic KCC2 (Fig 3B; Supplemental Fig 2B, C shows membrane and cytosolic values separately). For the cervical enlargement ECR motor pool, comparison of ipsilesional and contralesional labeling in both early and chronic timepoints show no difference in KCC2 ratios. Importantly, these ratios are not significantly different from the naïve, ruling out a bilateral reduction (see Fig 3 legend for statistics). Also, there was no change in lumbar KCC2 (see Fig 3C legend for statistics), which is expected since there was no RDD reduction in the contralesional hindlimb.

**Figure 3.**
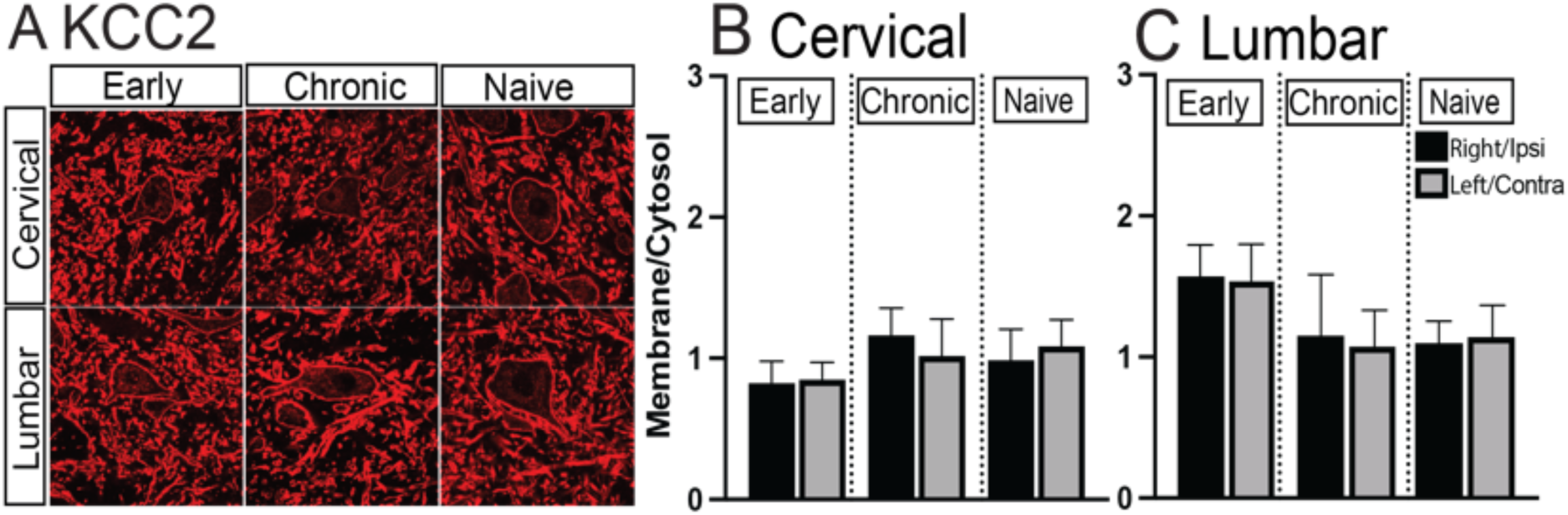
Motor neuron KCC2. A. Confocal micrographs. KCC2 immunostaining in the cervical and lumbar levels for the early and chronic cohorts, and naïve rats. B. Ratio of membrane/cytosolic KCC2 for cervical cord showed no significant difference between early, chronic, and naïve groups (one-way ANOVA; F=0.507; p=0.77). C. Same as B but for lumbar cord (one-way ANOVA; F=0.756; p=0.58).

To validate KCC2 immunohistochemistry, we performed western blot analysis and calculated relative protein levels of KCC2 from extracted cervical and lumbar tissue (Supplemental Figure 3). Analysis of ipsilesional, contralesional, and naive KCC2 from both cervical and lumbar levels collected at the early timepoint show no significant differences (Supp Fig 3 legend). Further, the absence of a KCC2 effect is not due to normalizing gray matter levels to white matter KCC2 (Boulenguez et al., 2010). Comparisons of ipsilesional, and contralesional sides, at early and chronic timepoints, and with the two sides in naïve rats, show no significant differences in the cervical membrane KCC2 (Supplemental Fig 1A; statistics described in legend). The same is true for comparisons done with labeling in the cervical cytosol, lumbar membrane, and lumbar cytosol (Supplemental Fig 1B, C, D; see legends for statistic values).

Taken together, our findings show that KCC2 is not a contributing mechanism driving motor neuronal hyperreflexia after unilateral PTX.

### VGlugt1 1A terminals increase on cervical and lumbar motor neurons but GABApre terminals increase only on lumbar motor neurons

Previously we showed that unilateral PTX produces afferent fiber sprouting as increased axon density in both the dorsal horn and intermediate zone, as well as increased 1a terminals on motor neurons (Tan et al., 2012). With hyperreflexia occurring only in the contralesional forelimb after PTX, and no reduction in membrane KCC2, we next examined 1a afferent circuit changes at contralateral cervical and lumbar levels. We used high-resolution 3-D reconstruction (Imaris; Figure 4A) to monitor all contacts between VGlut1+ afferent terminals on single motor neurons and their associated GABA presynaptic inhibitory synapses. In a representative control ECR motor neuron, we see VGlut1-labeled presynaptic terminals (red) scattered over the somatic and dendritic membrane (4A). GAD-positive GABApre terminals are located on many of the VGlut1 terminals (Betley et al., 2009). An enlargement of the boxed area (A, top right) and associated single optical slice (A, bottom right) reveal a cluster of synapses. After PTX, we observed in the cervical cord, more 1A terminals on the contralesional compared with ipsilesional cervical motor neurons (4B, red arrows), showing axon terminal sprouting. By contrast, the number of GABApre (green) on the two sides were similar. The contralesional lumbar motor neuron (4B, bottom) revealed both increased 1A presynaptic terminals (red arrowheads) and a commensurate increase in GABApre presynaptic terminals (green) on the 1A terminals compared with the ipsilesional side.

**Figure 4.**
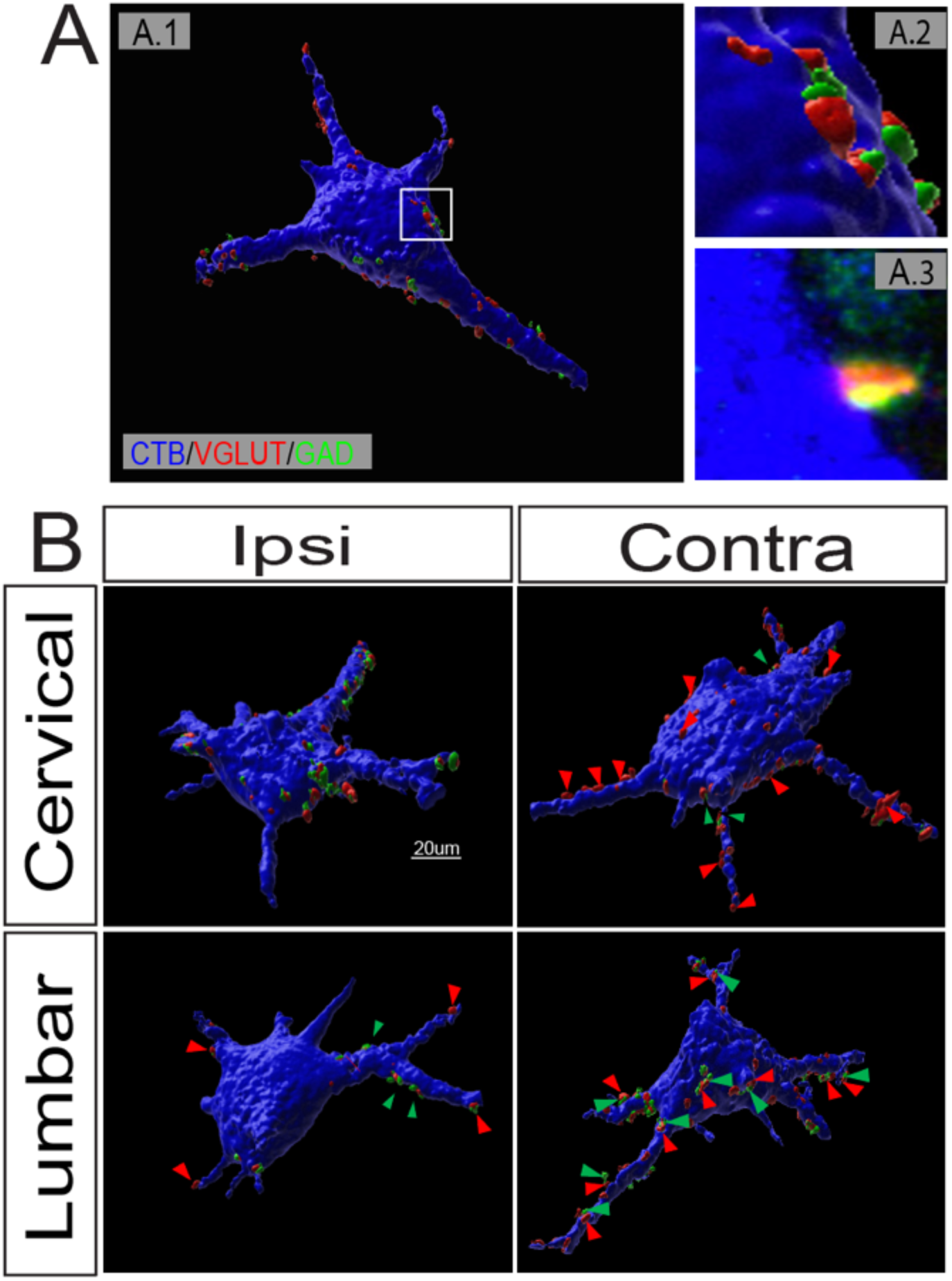
3-dimensional reconstruction of motor neurons. A. Representative motor neuron from a naïve rat (A1, whole-cell reconstruction; A2, 3-D cluster of synapses; A3, single optical slice). B. Representative motor neurons after unilateral PTX, at the early timepoint. (Calibration: 20µm).

Quantification of these morphological changes for the cervical segment (Figure 5A) revealed significant increases in 1A terminals, normalized to motor neuron surface area, for both the 7 dpi and 42 dpi time points (7 dpi: unpaired t-test 0.001. ± 0.0003; p<0.0005; 42 dpi: unpaired t-test test 0.001 ± 0.0003; p<0.0008; Fig 5A). We then quantified the number of GABApre at early and chronic timepoints and found no change between contralesional, and ipsilesional in the cervical spinal cord (Fig 5B).

**Figure 5.**
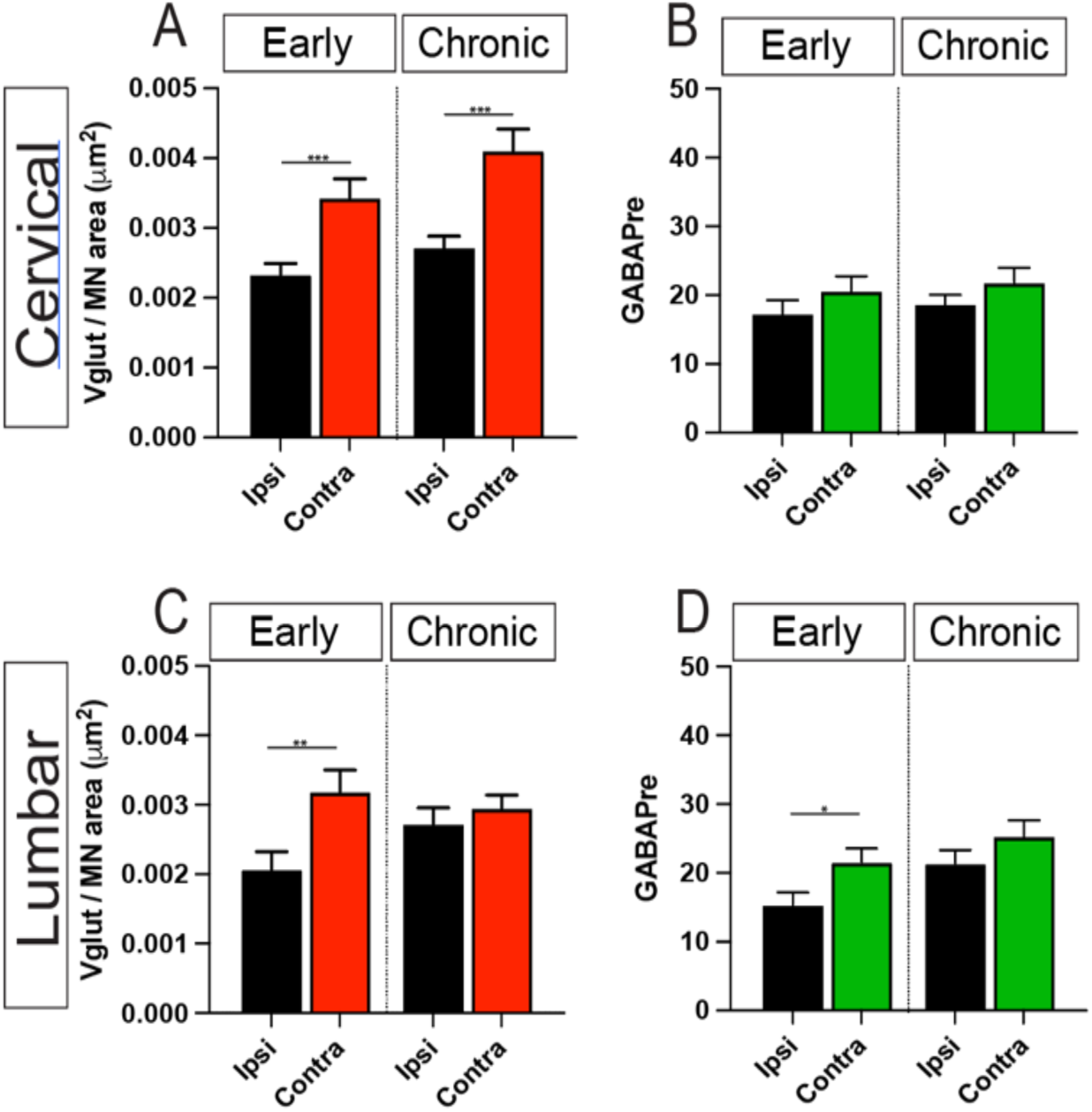
Quantification of 1A afferent and GABApre synapses after unilateral PTX; data from the early and chronic cohorts are shown. A. Cervical 1A synapses. B. Cervical GABApre (early-unpaired t-test; 1.663 ± 2.291; p<0.47), (chronic-unpaired t-test; 3.160 ± 2.550; p<0.21). C. Lumbar 1A synapses (unpaired t-test 0.0002288 ± 0.0002961; p=0.44) D. Lumbar GABApre (chronic-unpaired t-test; 3.971 ± 3.118; p<0.206).

In the lumbar cord, we observed a significant increase in normalized 1A terminals on TA motor neurons at the early (unpaired t-test 0.001 ± 0.0004; p<0.0083) (Fig5C). Importantly, unlike in the cervical cord at the early time point, we observed a significant increase in GABApre synaptic inputs (unpaired t-test 3.88 ± 2.94; p<0.014) (Fig. 5D) suggesting appropriate presynaptic inhibitory regulation of the actions of afferent sprouting. At the chronic timepoint, there was neither a significant increase in 1A terminals nor a significant increase in GABApre boutons. These findings show sprouting of 1A presynaptic terminals for both the cervical and lumbar motor neurons early after PTX, but a significant increase in GABApre terminals only in the lumbar cord. Interestingly, for the chronic timepoint, the lack of differences between contralesional and ipsilesional 1A and GABApre boutons suggests pruning from earlier afferent fiber sprouting, with commensurate GABApre increases.

## Discussion

In addition to upper limb muscle weakness and loss of voluntary control after injury to the motor systems, hyperreflexia is a key gain-of-function impairment that disrupts control of arm and hand movements. Hyperreflexia includes stronger reflexes with aberrant patterns of muscle contraction and spasticity (Thompson et al., 1992). In this study using unilateral CST loss, we found dissociations between 1A afferent fiber circuit plasticity and motor neuron membrane bound KCC2 that provide new insights into the mechanisms of hyperreflexia (Figure 6, which summarizes data for the contralesional side at early and chronic timepoints). Only the contralateral forelimb developed hyperreflexia, as measured by a reduction in RDD. This leads to the expectation that an injury-dependent change associated with, and possibly causal to, hyperreflexia in this model will be present in the contralateral cervical cord only. KCC2 reduction, key to hyperreflexia after SCI and bilateral CST lesion (Kathe et al., 2016; Chen et al., 2018; Beverungen et al., 2020), does not associate with cervical hyperreflexia in the unilateral PTX model, showing that hyperreflexia can occur without enhanced motor neuronal excitability due to downregulation of KCC2. Whereas 1A sprouting is similarly present at both the cervical and lumbar levels at 7dpi, likely due to loss of synaptic competition after CST lesion (Jiang et al., 2016; Jiang et al., 2019), hyperreflexia is only present in the forelimb.

**Figure 6.**
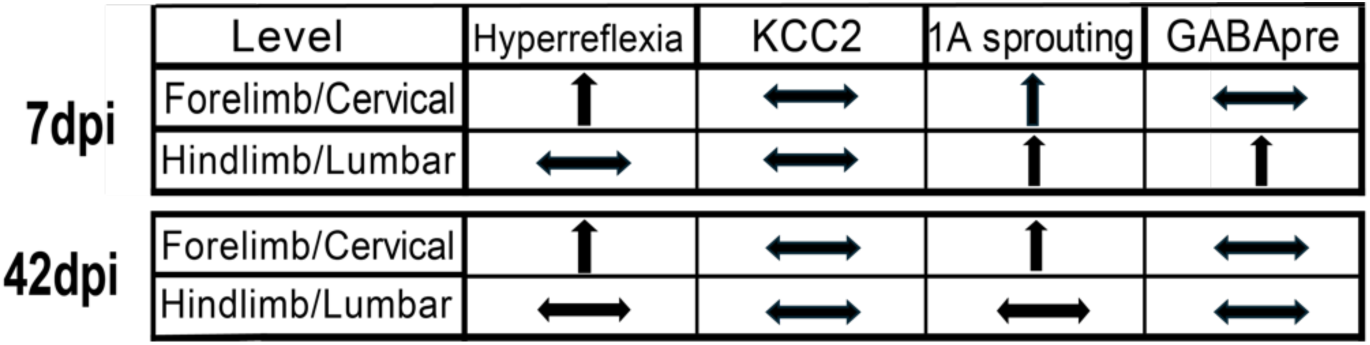
Summary of contralesional forelimb and hindlimb changes in reflex strength, measured as rate-dependent depression, and corresponding changes in KCC2, 1A afferent density on motor neurons, and GABApre at the cervical and lumbar levels, at 7-dpi and 42-dpi. Note, there were no changes ipsilesionally.

Thus, 1A sprouting alone does not associate with hyperreflexia when the cervical and lumbar segments are compared. Finally, the lack of a change in GABApre in the cervical cord suggests that 1A afferent sprouting leads to enhanced reflex signaling because of unchecked presynaptic inhibitory regulation of the sprouted 1A terminals. By contrast, the increase in GABApre terminals in the lumbar spinal cord parallels the increase in 1A terminals. This is consistent with effective presynaptic inhibitory regulation for hindlimb 1A afferent signaling and, in consequence, no hyperreflexia.

### The forelimb is more vulnerable than the hindlimb to developing hyperreflexia

Clinical studies show that after unilateral cortical stroke in humans, spasticity is more prevalent and more severe in the upper limb than for the lower limb (Katoozian et al., 2018; Sunnerhagen et al., 2019). Paralleling this finding, we found that unilateral PTX produces forelimb not hindlimb hyperreflexia. As described above, we have used this finding in support of potential causality of a lack of a compensatory increase in GABApre terminals failing to check 1A afferent sprouting, and contributing to hyperreflexia in the unilateral PTX model. This is a novel finding, as animal studies examine the forelimb or hindlimb, but not both. Furthermore, animal studies suggest that the CST may have differential cervical and lumbar projection densities and functional roles for movement control of the forelimb and hindlimb. Whereas direct comparisons of CST cervical and lumbar CST projections are limited, animal studies show that forelimb MCX projects to the cervical cord, but hindlimb MCX projects to both cervical and lumbar segments (Steward et al., 2021), pointing to a stronger CST role in forelimb control.

Classical tracing studies (Armand et al., 1985), confirmed by genetic tracing (Bareyre et al., 2005), show the typical reduction in CST white matter axon numbers from rostral to caudal, which is largely due to gray matter axon termination. But accompanying the rostro-caudal tract reduction, there is also a profound reduction in gray matter terminations within the thoracic and lumbar segments (Armand et al., 1985). Denser CST projection to the cervical than lumbar gray matter in the intact spinal cord may underpin the differential vulnerability of the fore-and hindlimb, and associated spinal circuits, to hyperreflexia after unilateral CST loss. Hyperreflexia in this model may thus reflect a ‘dose effect.’ Intriguingly, with CST presynaptic contacts as likely key determinants of the postsynaptic response to injury, providing a surrogate source of the activity they contributed (e.g., rehabilitation or neuromodulation) may facilitate recovery after SCI or stroke.

### Hyperreflexia presents acutely after injury and is maintained into the chronic period

Our assessment of hyperreflexia showed RDD was numerically reduced at day 2, the earliest time point tested, and was significant at 1-week post-injury. This early presentation suggests a pathophysiological process triggered by the sudden loss of connections (Jiang et al., 2016) or their activity (Jiang et al., 2019). The significant RDD reduction at 1-week post-injury was not different from 6-weeks, showing persistence of hyperreflexia. This early presence and maintenance over several weeks, led us to consider the process of hyperreflexia development. The RDD reduction at the earliest timepoint, although not significant, suggests an immediate circuit level reorganization: the sudden loss of CST connections led to competitive primary afferent sprouting after injury (Jiang et al., 2016). The rapid onset and lack of progressive worsening over the 6 weeks of testing parallels observations in humans after stroke. Upper limb spasticity occurs 3 days after first presentation of a stroke in 25% of patients examined (Opheim et al., 2014; Opheim et al., 2015). Using logistic regression analysis, the best predictor for persistent spasticity at 12 months was spasticity at 10 days post injury (Sunnerhagen et al., 2019).

We identified sprouting of glutamatergic terminals on motor neurons at 1-week post injury, showing that a morphological change can occur remarkably early. Additionally, in the lumbar cord, GABApre terminals also showed an increase at this early time point. Unilateral photothrombotic motor cortex stroke in the mouse produces an early reduction in RDD (3-days) and persistence into the chronic period (7-weeks) (Toda et al., 2014), although the magnitude of the reduction is variable (Toda et al., 2014; Lee et al., 2014). Similarities between unilateral selective CST injury and stroke, as they relate to time course of hyperreflexia in animals and spasticity onset in humans, further establish the unilateral PTX model as an important tool to inform therapeutic strategies aimed at reducing spasticity after a brain injury.

Constitutive activation of the serotonin receptor 5-HT_2c_ is another well studied mechanism contributing to motor neuron hyperexcitability and spasticity in humans and rodents after SCI (Gorassini et al., 2004; Murray et al., 2010). The switch to constitutive 5-HT_2c_ activation following spinal cord injury is thought to be provoked by the loss of descending monoaminergic inputs to the spinal cord. Although the shift to constitutively active 5-HT_2c_ was not examined in this study, it would not be expected to occur after selective CST loss.

Furthermore, this mechanism is not expected to contribute to hyperreflexia early on. Study of the time course of 5-HT_2c_ receptor expression in rats after SCI estimates 1.3x fold increase in 5-HT_2c_ receptor activity when compared to the sham group as early as 14 days after injury (Ren et al., 2013). This time course indicates that the shift to the constitutively active form cannot explain our finding that hyperreflexia begins within days after PTX. Although not significantly different with longer post-injury survival periods, RDD seems to be reducing further post-injury. This may reflect a delayed action of constitutive 5-HT_2c_ activity (Ren et al., 2013).

### Do spared ipsilateral CST axons protect from increased excitability after unilateral PTX?

We found no evidence for a reduction in membrane-bound KCC2 contralesionally. This shows that hyperreflexia after unilateral CST loss is not due to KCC2 mediated motor neuron hyperexcitability. SCI and bilateral PTX, which completely lesion all CST fibers have identified KCC2 downregulation, as an important contributor to enhancing motor neuron excitability and, in turn, hyperreflexia (Boulenguez et al., 2010; Kathe et al., 2016; Beverungen et al., 2020).

Comparing immunohistochemical findings for the affected and unaffected sides after unilateral motor cortex stroke in the mouse suggest transient and bilateral reductions in KCC2, which were not corroborated using western blotting (Toda et al., 2014). By contrast, we show no changes either with immunohistochemistry or western blotting. Therapeutic interventions targeting restoration of KCC2 are associated with a significant reduction of hyperreflexia, and spasticity (Murray et al, 2010; Kathe et al., 2016; Bilchak et al., 2021). Though the association between these injury models and KCC2 downregulation is clear, the unilateral PTX model used in this study shows evidence of a disassociation between CST injury, hyperreflexia, and motor neuron hyperexcitability. This knowledge informs therapy after unilateral injury (i.e., motor cortex stroke) in that KCC2 support would not be expected to improve function as an activity-based approach that could reshape the 1A afferent circuit.

The question remains why with unilateral PTX, KCC2 is not involved when it is a contributing factor after bilateral CST lesion and SCI. In our hands, we also observe a significant reduction in KCC2 after bilateral PTX (unpublished observations). The presence of ipsilateral CST projections may be sufficient to protect against hyperreflexia by providing necessary segmental excitatory input; bilateral CST lesion eliminates that protection. Moreover, there is evidence of injury-induced plasticity of spared ipsilateral CST axons after a unilateral injury; whereby, with the loss of the contralateral CST, ipsilateral uninjured CST fibers sprout (Bareyre et al., 2005; Brus-Ramer et al., 2007) and may even contact motor neurons (Bareyre et al., 2005; Weidner and Tuszynski, 2002). There can be a doubling of this injury-dependent ipsilateral CST sprouting (Brus-Ramer et al., 2007), which could provide compensatory connections to help support segmental excitability. The support provided by these spared CST projections could be further strengthened with electrical stimulation (Brus-Ramer et al., 2007). It is possible the ipsilateral CST projects sufficient excitatory inputs to mitigate motor neuron hyperexcitability that results from KCC2 reduction, but not enough to prevent hyperreflexia in the affected forelimb.

As emphasized earlier, hyperreflexia can be mediated by multiple factors. The unilateral PTX model demonstrates a dissociation. Among the three factors that we have examined—1A sprouting, reduced GABApre regulation, and reduced membrane-bound KCC2—each could, in principle, have led to a stronger reflex. But only the persistent lack of cervical GABApre sprouting, along with 1A afferent sprouting, explained restricted hyperreflexia in the contralateral forelimb. Considering our findings in the broader context of motor system injuries, our findings demonstrate a circuit disorder, without need for KCC2 motor neuron hyperexcitability, mediating hyperreflexia. The physiological actions of sprouting of 1A afferent glutamatergic terminals on motor neurons are not checked by concomitant GABApre sprouting; there is a higher proportion of VGlut1 terminals on motor neurons without GABApre regulation.

## Acknowledgements

We thank XiuLi Wu for histology. The authors have no competing interests to declare.

## Supplemental Figures

**Supplemental Figure 1.**
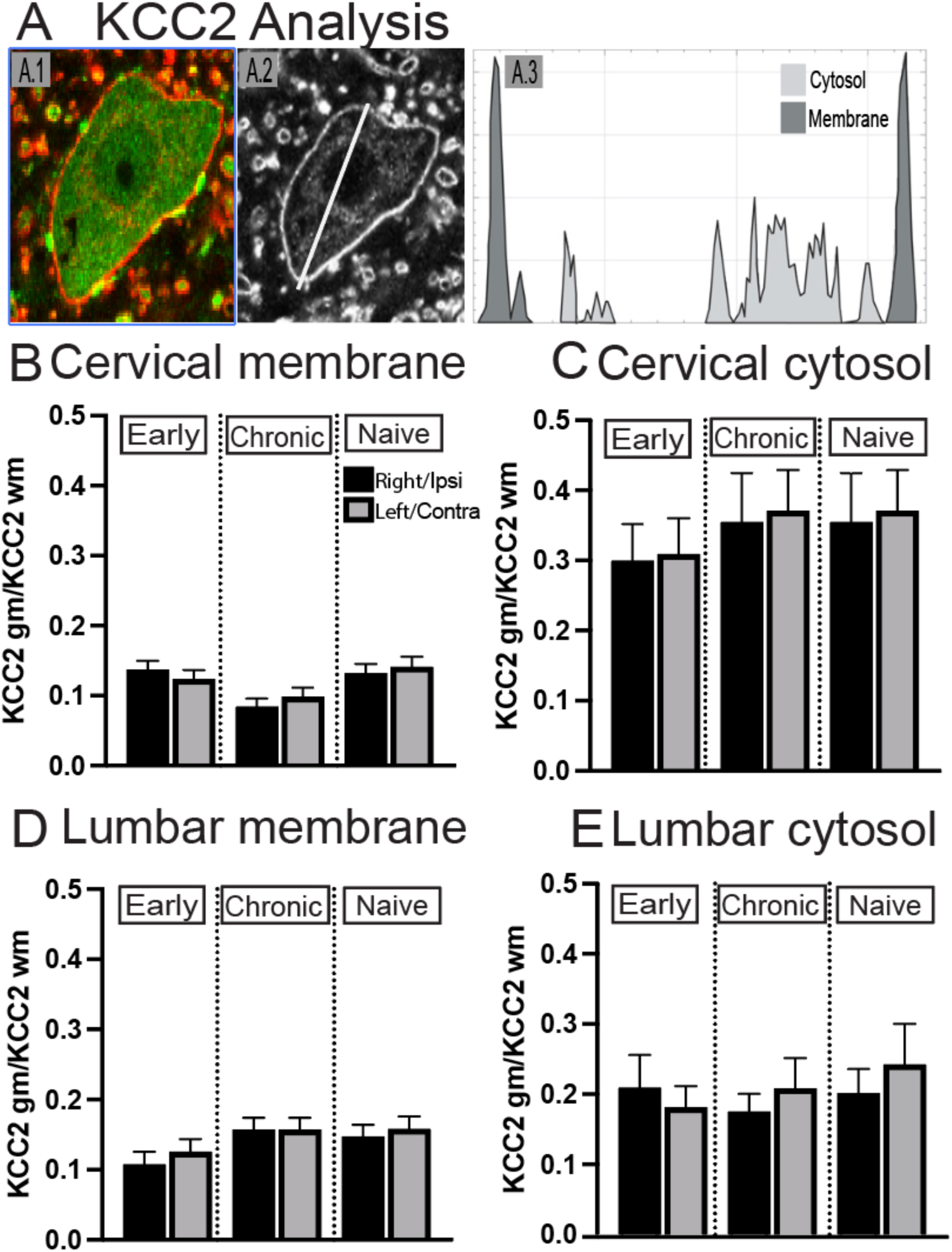
KCC2 immunohistochemistry measurement and analyses. KCC2 Normalized membrane/cytosol averages. A. Methods. A1. Optical slice of a representative single motor neuron. Red, KCC2; Green ChAT. A2. Example of drawn line to measure KCC2 levels. A3. ImageJ plot showing pixel level along the line, showing membrane (dark gray) and cytosol (light gray) values. B. Cervical membrane comparison (one-way ANOVA; F=2.030; p=0.079). C. Cervical cytosol comparison (one-way ANOVA; F=0.519; p=0.760). D. Lumbar membrane comparison (one-way ANOVA; F=1.872; p=0.141). E) Lumbar cyt[osol comparison (one-way ANOVA; F=0.234; p=0.872).

**Supplemental Figure 2.**
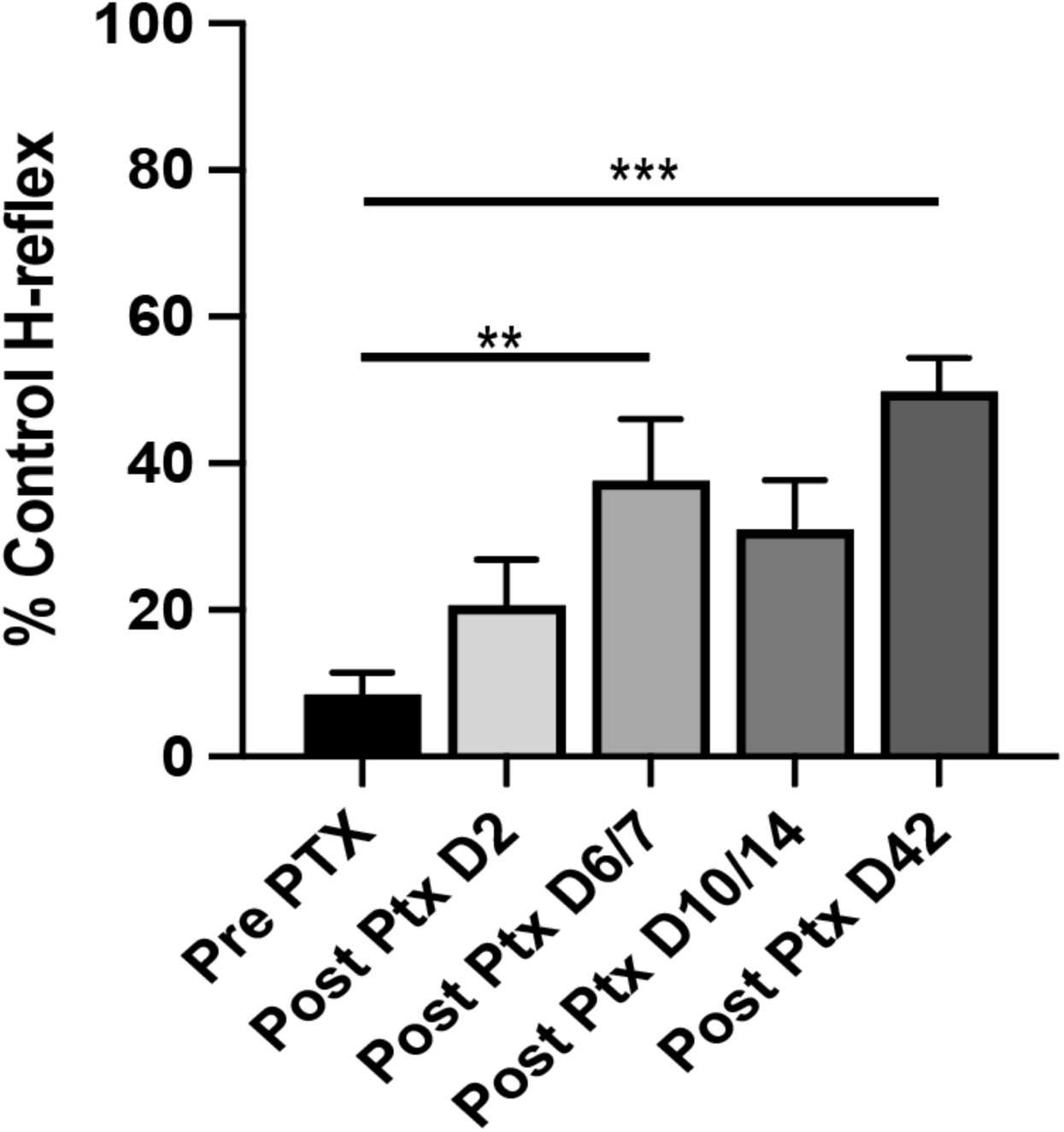
Time course of hyperreflexia development. Multiple comparison analysis based on main, and supplemental animal cohorts (n=11) shows the numerical increase in hyperreflexia (i.e., increase in percent control measure, RDD at 20 ms) from 2-dpi to 6/7 dpi with apparent stabilization of values thereafter. (6/7-DPI; one-way ANOVA; F=6.548; p=0.005; 42-dpi, one-way ANOVA; F=6.548; p=0.0005).

**Supplemental Figure 3.**
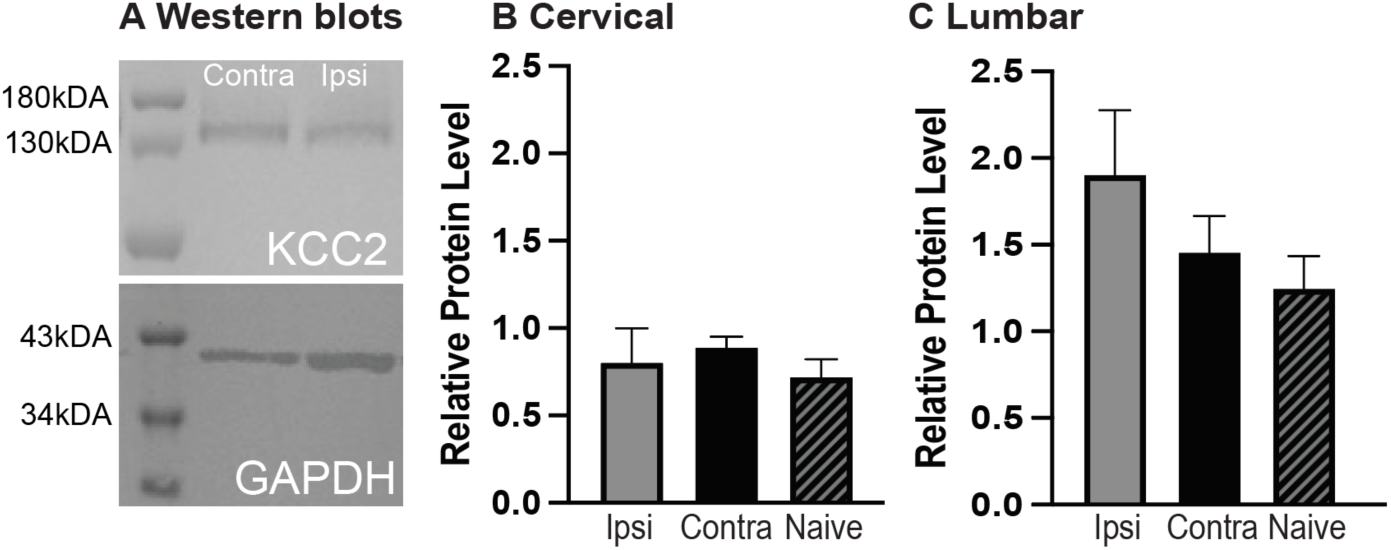
Western blotting for KCC2. A. Representative western blots from cervical (top) and lumbar (bottom) spinal tissue. B. Cervical protein level comparisons (one-way ANOVA; F=0.4697; p=0.63). C. Lumbar protein level comparisons (one-way ANOVA; F=1.789; p=0.19).

**Supplemental Figure 4.**
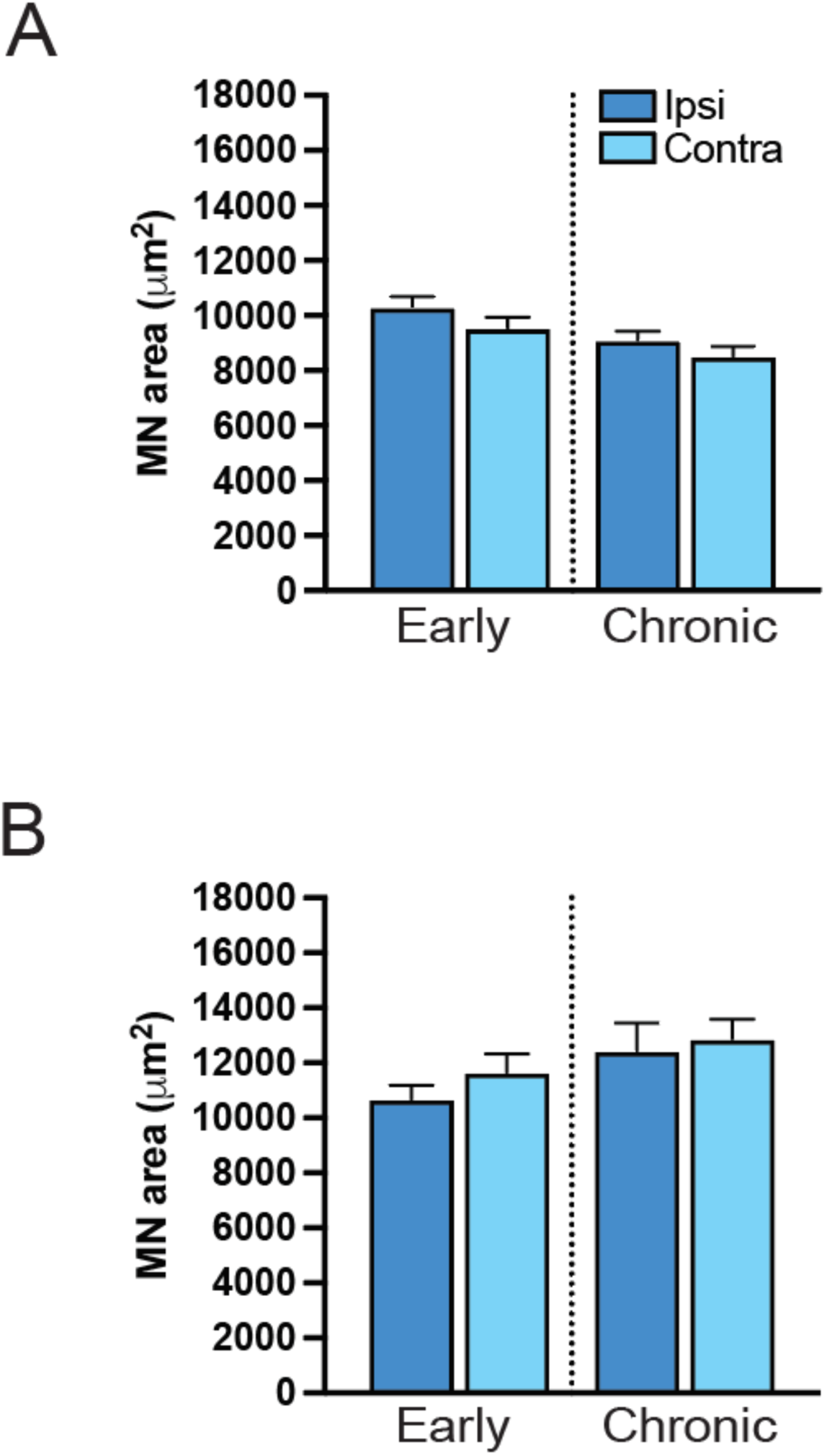
Motor neuron surface area averages. Quantification of average ipsilateral, and contralateral motor neuron size by surface area (µm^2^). A. Cervical, early and chronic comparisons (one-way ANOVA; F=1.71; p=0.16). B. Lumbar, early and chronic comparisons (one-way ANOVA; F=1.68; p=0.17).

**Supplemental Figure 5.**
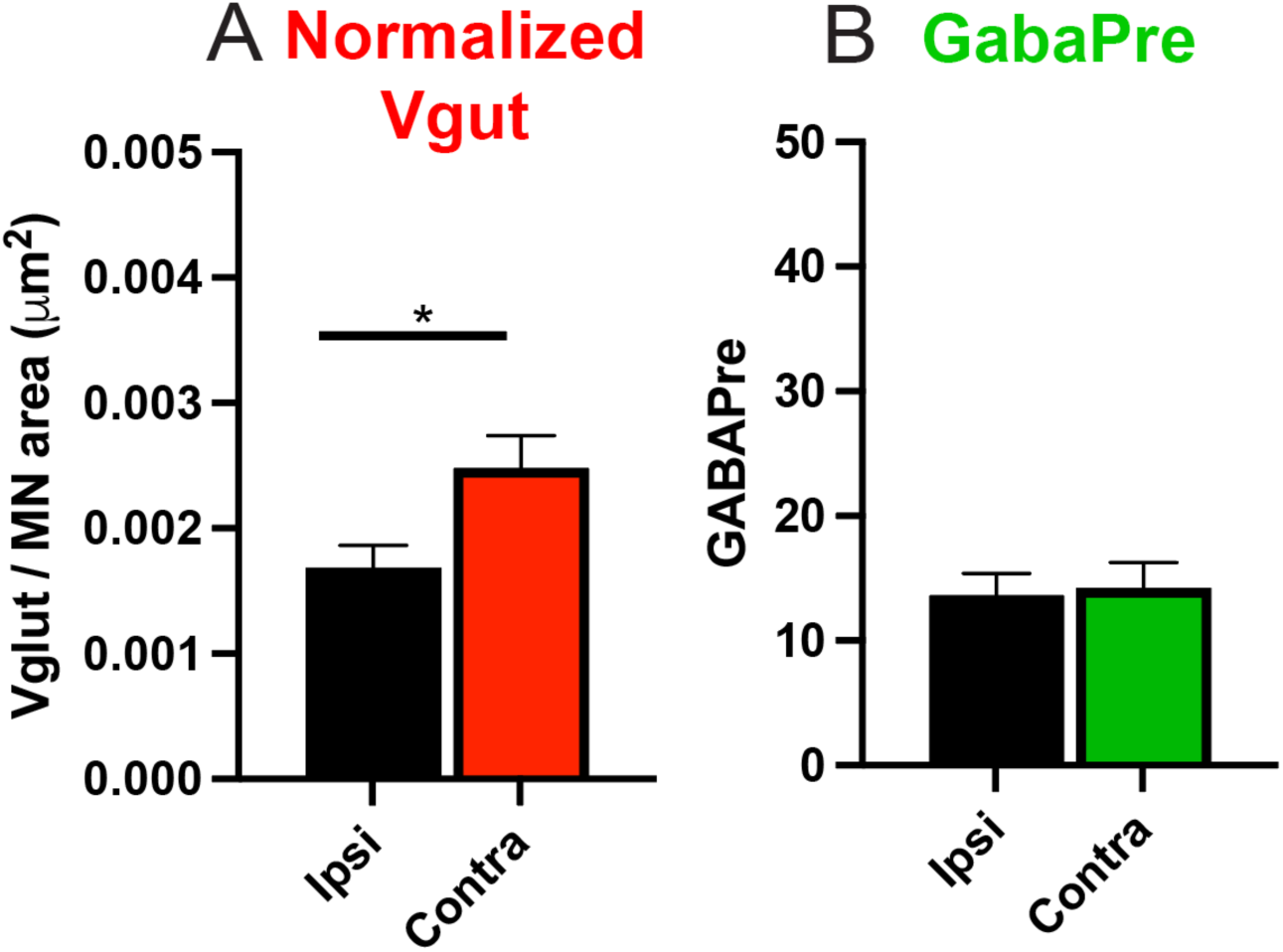
Cervical level assessment of normalized VGlut1 and GABApre-positive contacts on ADQ motor neurons. A. Normalized VGlut1 (unpaired t-test 0.0007970 ± 0.0003057; p=0.011). B) GABApre (unpaired t-test 0.6022 ± 2.694; p=0.823).

