## Supplemental Figures for "Hyperreflexia after corticospinal tract lesion reflects 1A afferent circuit changes not increased KCC2 hyperexcitability"

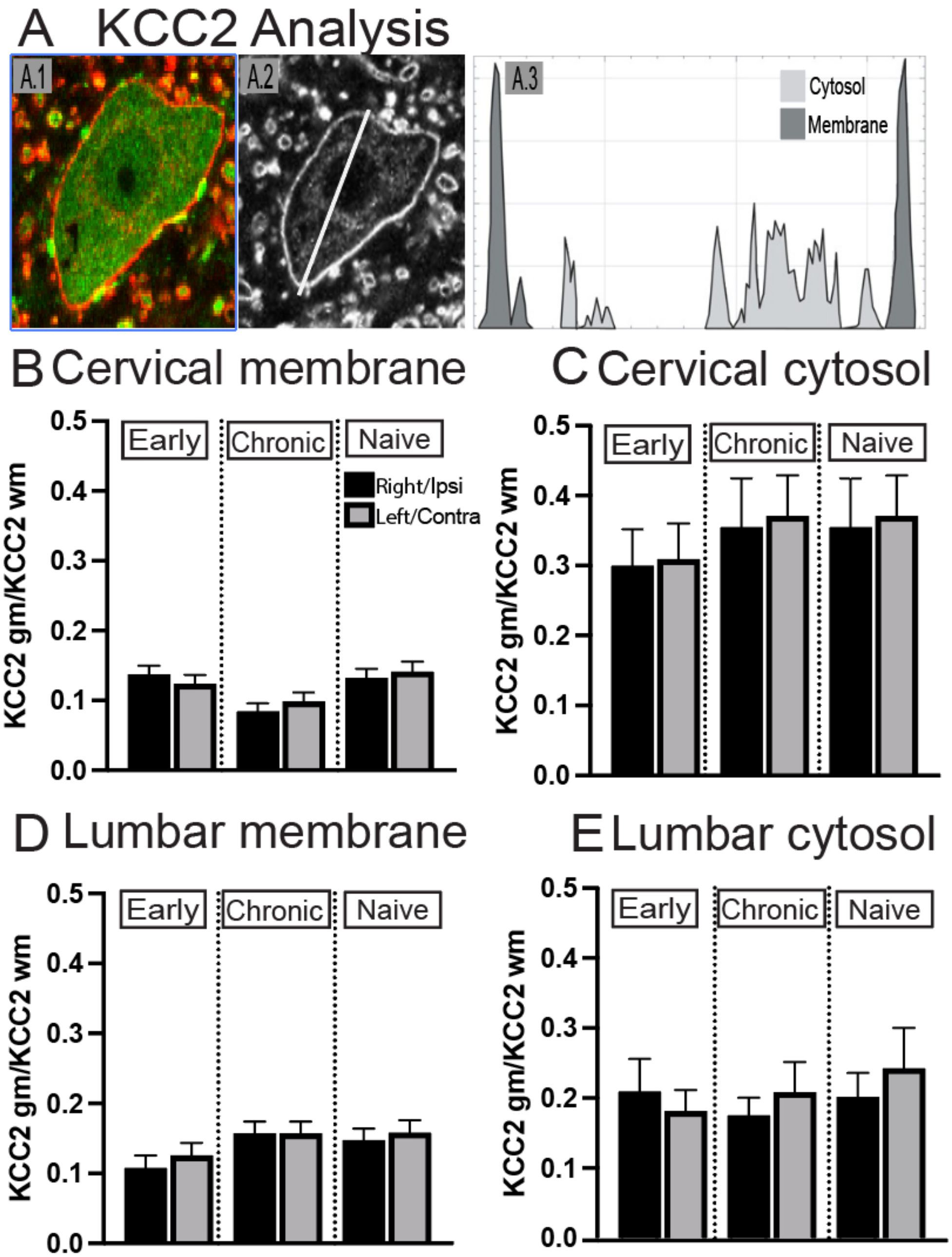


**Supplemental Figure 1**. KCC2 immunohistochemistry measurement and analyses. KCC2 Normalized membrane/cytosol averages. A. Methods. A1. Optical slice of a representative single motor neuron. Red, KCC2; Green ChAT. A2. Example of drawn line to measure KCC2 levels. A3. ImageJ plot showing pixel level along the line, showing membrane (dark gray) and cytosol (light gray) values. B. Cervical membrane comparison (one-way ANOVA; F=2.030; p=0.079). C. Cervical cytosol comparison (one-way ANOVA; F=0.519; p=0.760). D. Lumbar membrane comparison (one-way ANOVA; F=1.872; p=0.141). E) Lumbar cyt[osol comparison (one-way ANOVA; F=0.234; p=0.872).

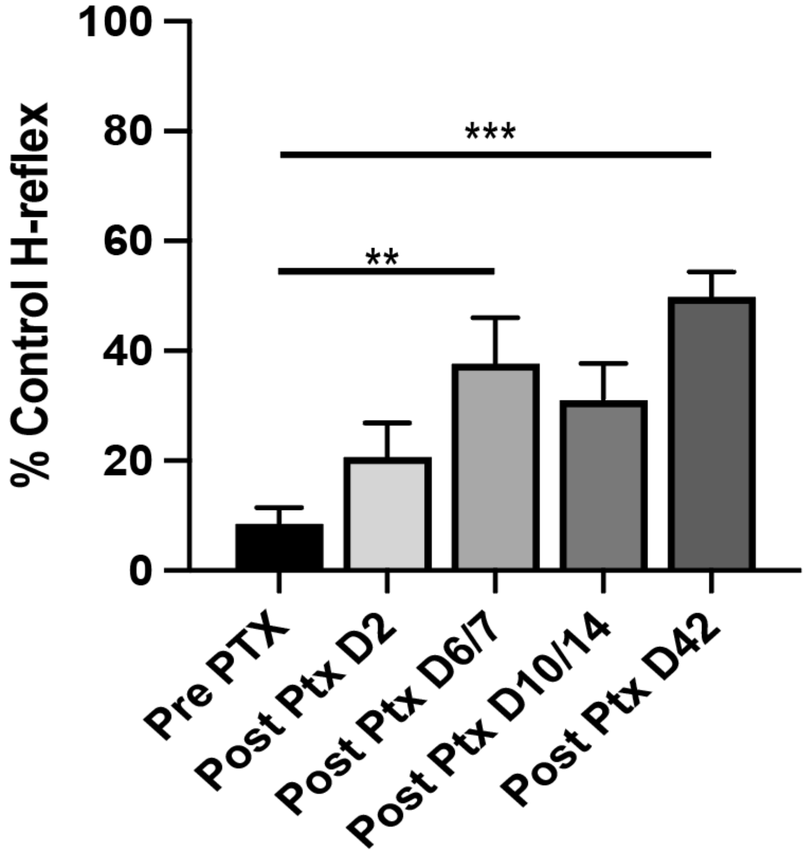


**Supplemental Figure 2.** Time course of hyperreflexia development. Multiple comparison analysis based on main, and supplemental animal cohorts (n=11) shows the numerical increase in hyperreflexia (i.e., increase in percent control measure, RDD at 20 ms) from 2-dpi to 6/7 dpi with apparent stabilization of values thereafter. (6/7-DPI; one-way ANOVA; F=6.548; p=0.005; 42-dpi, one-way ANOVA; F=6.548; p=0.0005).

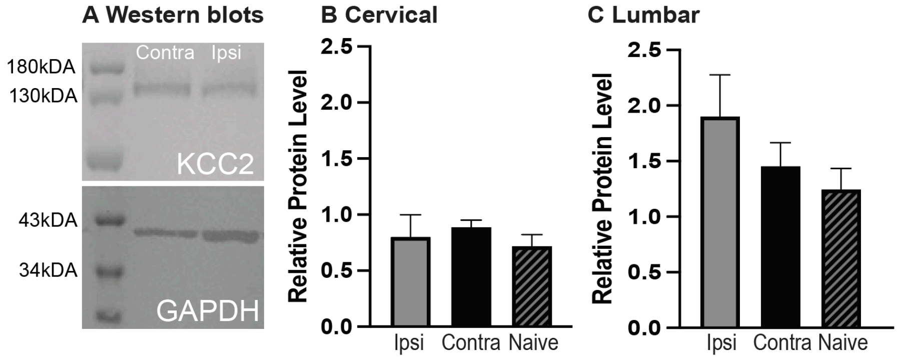


**Supplemental Figure 3**. Western blotting for KCC2. A. Representative western blots from cervical (top) and lumbar (bottom) spinal tissue. B. Cervical protein level comparisons (one-way ANOVA; F=0.4697; p=0.63). C. Lumbar protein level comparisons (one-way ANOVA; F=1.789; p=0.19).

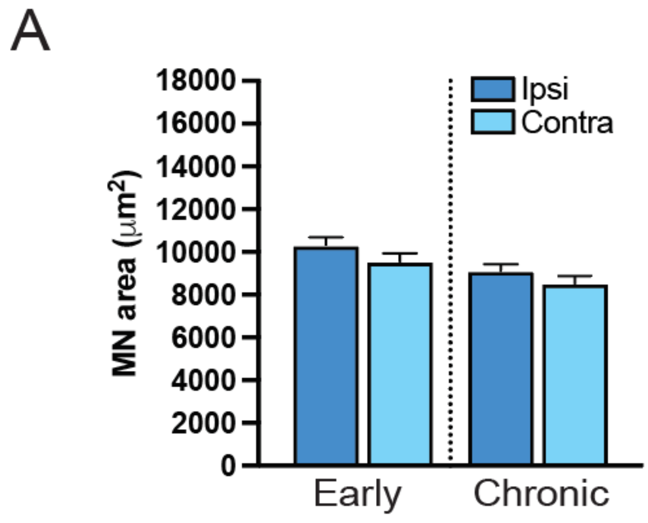


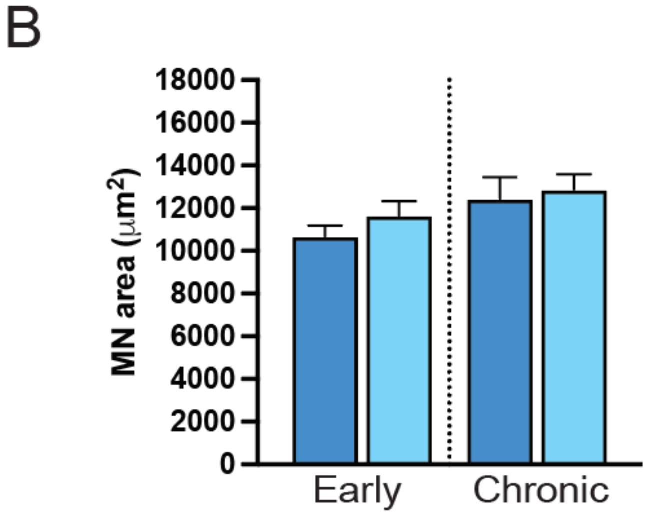


**Supplemental Figure 4.** Motor neuron surface area averages. Quantification of average ipsilateral, and contralateral motor neuron size by surface area (μm$^2$). A. Cervical, early and chronic comparisons (one-way ANOVA; $F=1.71$; $p=0.16$). B. Lumbar, early and chronic comparisons (one-way ANOVA; $F=1.68$; $p=0.17$).

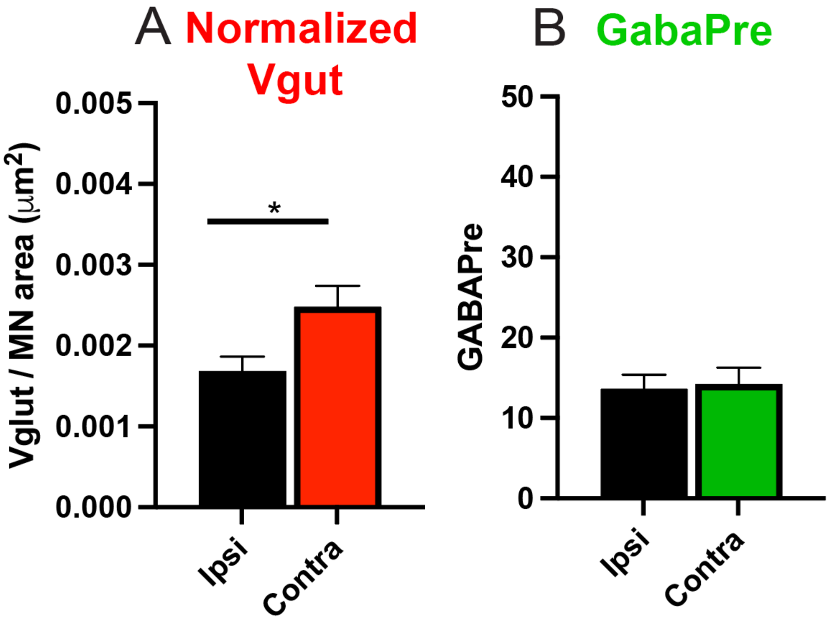


**Supplemental Figure 5**. Cervical level assessment of normalized VGlut1 and GABApre-positive contacts on ADQ motor neurons. A. Normalized VGlut1 (unpaired t-test 0.0007970 ± 0.0003057; p=0.011). B) GABApre (unpaired t-test 0.6022 ± 2.694; p=0.823).
